# Single-cell Landscape of Immune Cells in Blood and Skin in Psoriasis

**DOI:** 10.1101/2024.09.17.613463

**Authors:** Jingwen Deng, Michel Olde Nordkamp, Shuyan Ye, Jingjie Yu, Deepak Balak, Wanlin Yu, Timothy Radstake, José A. M. Borghans, Chuanjian Lu, Aridaman Pandit, Bram Gerritsen

## Abstract

**Background:** Psoriasis is a systemic inflammatory disease for which there is currently no cure, in part due to an incomplete understanding of its pathophysiology.

**Methods:** To better understand the immune response in psoriasis, we performed single-cell RNA sequencing (scRNA-seq) on peripheral blood mononuclear cells (PBMCs) and on lesional and non-lesional skin samples from a cohort of 11 psoriasis patients and 8 healthy controls. Additionally, we conducted flow cytometry on PBMCs from a separate cohort of 13 psoriasis patients and 11 ankylosing spondylitis.

**Findings:** Our study revealed altered immune signatures of specific myeloid and lymphocyte subsets in blood and skin, both in terms of cell numbers and gene expression. Specifically, we discovered elevated proportions of circulating CD14^++^ monocytes, increased expression of major histocompatibility complex (MHC) class II molecule by circulating CD16^+^ monocytes, as well as increased expression of genes related to skin homing and to pro-inflammatory responses in psoriasis by circulating plasmacytoid dendritic cells (pDCs). Circulating CD8^+^ T effector memory cells in psoriasis patients exhibited reduced abundance but increased skin-homing potential. In psoriatic lesions, we observed a hyperinflammatory myeloid-cell state and enrichment of IL17-producing cells with a tissue-resident memory T-cell signature.

**Interpretation:** The changes in immune cell numbers and gene expression indicate a significant alteration in the immune landscape of psoriasis patients. This suggests that the immune system in psoriasis is reprogrammed, affecting both innate and adaptive branches. These findings provide new insights into the aberrant immune-cell signatures in the circulation and skin lesions in psoriasis, and thereby help to understand its pathophysiology.

**Funding:** This study was financially supported by the National Natural Science Foundation of China (U23A6012), Science and Technology Planning Project of Guangzhou (2024A03J0055, 202206080005), Innovation Team and Talents Cultivation Program of National Administration of Traditional Chinese Medicine (ZYYCXTD-C-202204).

## Introduction

Psoriasis is a systemic inflammatory disease characterized by erythematous, scaly and pruritic skin plaques. The inflammation in psoriasis is not limited to the skin, and can also affect other organ systems, such as the joints, the cardiovascular-metabolic system, and the central nervous system. The pathophysiology of psoriasis is at least in part caused by abnormal keratinocyte proliferation, resulting from complex interactions between the innate and the adaptive immune system. In psoriatic skin, the crosstalk between T cells, dendritic cells (DCs), and keratinocytes is mainly mediated via their secreted cytokines. Accumulating evidence indicates that the IL-23/Th17 axis is the major immune pathway underlying the epidermal hyperplasia and immune cell infiltration in psoriasis [1]. IL-23 and IL-12 released by inflammatory myeloid DCs activate Th17 cells, Th22 cells and Th1 cells to produce pro-inflammatory cytokines, such as IL-17, IL-22, IFN-γ, and TNF. These cytokines act on keratinocytes, and thereby intensify psoriatic inflammation [2].

With our increased knowledge of disease pathogenesis, the number of available, effective targeted treatment options has risen significantly. Monoclonal antibodies directed against specific cytokines, such as anti-TNF, anti-IL-17 and anti-IL-23 biologics, are highly effective in treating psoriasis, with more than 70% of patients achieving a 75% or greater reduction in the psoriasis area and severity index (PASI) after treatment[3]. Yet, there are still some important obstacles to achieving a cure for psoriasis patients, such as a lack of response to treatment in at least 10% of patients [4], and infectious adverse events related to interleukin-antagonism, such as mucocutaneous candidiasis associated with IL-17-blockade [5]. Additionally, at least 30% of patients experienced a relapse after the discontinuation of biologics [6]. To better manage this complex condition, it is important to further our understanding of the immune dysfunction and the interaction between systemic and cutaneous immunity in psoriasis.

Single-cell RNA sequencing (scRNA-seq) has added a new dimension to our understanding of immune cell heterogeneity and diversity by providing detailed information about individual cells. This technology has been used to study psoriasis, and has revealed that multiple immune cell types, including myeloid cells, lymphoid cells, keratinocytes, and stromal cells are involved in the disease [7–11]. However, some studies lack validation of their findings. Nevertheless, simultaneous single-cell transcriptomic studies of immune cells in psoriasis in both circulation and skin are rare.

In this study, we applied single-cell transcriptome analysis to link data from immune cells in the circulation and the skin, in order to obtain a more detailed and informative view of the pathophysiology of psoriasis. We further validated our findings from single-cell analysis using an independent cohort through flow cytometry and public bulk transcriptome datasets (Figure.1A, B). We found that psoriasis is associated with systemic disruptions in the immune system, including changes in immune cell proportions and gene expression levels, with both specific subsets of the innate immune system (CD14^+^ monocytes, CD16^+^ monocytes and plasmacytoid dendritic cells (pDCs)) and subsets of the adaptive immune system (CD8^+^ effector memory T (T_EM_) cells and CD8^+^ tissue-resident memory T (T_RM_) cells) involved. These findings provide new insights into the pathogenesis of psoriasis and help explain the lack of consistent efficacy of biologics targeting a single molecule or pathway in psoriasis treatment.

**Figure 1.**
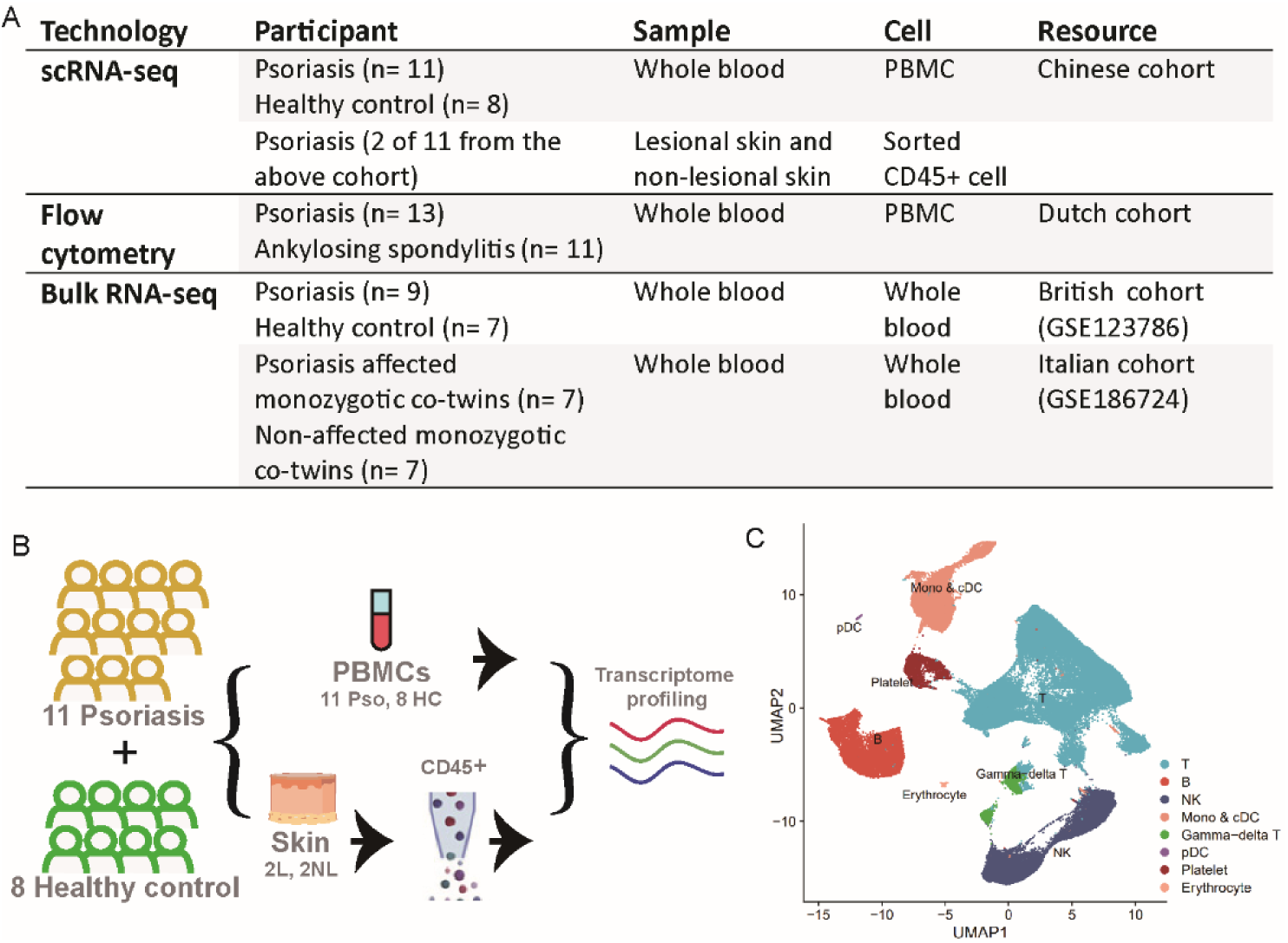
Overview of samples used and experimental design. A. Samples used in each of the experimental techniques. B. Schematic overview of the sample preparations and single-cell sequencing. C. UMAP visualization of PBMCs of 11 patients and 8 controls. The annotations for cell clusters are based on the DEGs of clusters (Figure.S1C) and the expression of marker genes (Figure.S1D).

## Materials and Methods

### Ethics statement and sample acquisition

scRNA-seq study was approved by the Institutional Ethics Committee of the Guangdong Provincial Hospital of Traditional Chinese Medicine (GPHCM B2020-238-02). Eleven patients with psoriasis were recruited, along with eight healthy donors who were matched for age and sex. After written informed consent had been given, all participants donated blood and two patients donated both skin biopsies and blood. None of the participants had applied biologics treatment before the start of the study, and all had been free of any other systemic treatment for at least 4 weeks and free of topical corticosteroids for at least 2 weeks. Participant recruitment took place at the outpatient clinic of the Department of Dermatology at Guangdong Provincial Hospital of Traditional Chinese Medicine from December 2020 to February 2021. The information of all participants is listed in supplementary Table 1.

Flow cytometry analysis was carried out at the University Medical Centre Utrecht (UMCU) in accordance with the Helsinki principles. Peripheral blood mononuclear cells (PBMCs) samples and clinical data were collected from a cohort of 13 psoriasis patients and 12 ankylosing spondylitis (AS) patients, who met the Assessment of SpondyloArthritis International Society (ASAS) criteria, in a prospective observational study. The AS patients served as a non-psoriasis control group, with none having a history of psoriasis. Participant recruitment took place at the outpatient clinic of the Department of Rheumatology and Clinical Immunology at UMCU from March 2016 to September 2018. The information of all participants is listed in supplementary Table 2.

### Cell isolation

Donors provided whole blood samples collected in EDTA-containing tubes. The blood was then diluted with PBS and separated using Ficoll density gradient medium. The PBMC layer was carefully collected using a sterile pipette and transferred to a new centrifuge tube. PBMCs were subsequently washed with PBS and utilized for library preparation.

Skin biopsies were collected from both lesional and non-lesional skin areas on the legs, with non-lesional areas being selected at least 2 cm away from the lesions. Fresh skin samples were trimmed to remove hair and subcutaneous adipose tissue, finely minced, and then incubated in 5 mL of RPMI containing 0.25 mg/mL Liberase and 1 mg/mL DNase for 2 hours at 37℃. In the last 15 minutes of the incubation period, 1 mL of 0.25% Trypsin-1 mM EDTA was added to the suspension. The digestion was terminated by adding 5 mL of 4℃ 5% FACS buffer. Cell suspensions were obtained after final dissociation using a 10 mL syringe, after which they were filtered, counted, and sorted for CD45^+^ cells.

### Flow cytometric analysis

Immune phenotype characterization of PBMCs sample was performed by flow cytometry, employing both surface and intracellular staining methods. Cell suspensions (1×10⁶ cells/ml) were stimulated for 3 hours at 37°C with PMA (25 ng/ml) and ionomycin (1 µg/ml) in the presence of Brefeldin-A (10 µg/ml). Prior to staining, cells were resuspended at a concentration of 1×10⁶ cells/100 µl per well in a 96-well plate, and blocked with 2 µg/ml of anti-CD16/CD32 diluted in PBS with 1% FBS, incubated at 4°C for 15 minutes to prevent non-specific Fc receptor binding.

Surface staining was conducted by incubating the cells in the dark at 4°C for 30 minutes using the following antibodies: Fixable Viability Dye eFluor™ 780, Hu CD3 Alexa Fluor® 700, Hu CD19 Alexa Fluor® 700, and Hu CD56 Alexa Fluor® 700. For the monocyte panel, Hu CD141 BV711, Hu CD123 FITC, Hu CD14 PerCP-Cy™5.5, Hu CD16 BV510, and Hu HLA-DR BV421 were used, while the T cell panel included Hu CD4 BV711, Hu CD8 BV500, Hu CCR4 BV605, and Hu CLA Pacific Blue™. Intracellular staining for the monocyte panel was performed using Hu TNF-alpha PE-Cy7.

The stained cells were analyzed using a BD LSRFortessa flow cytometer, and data were processed with FlowJo software (Version 10.0.8, Tree Star, Ashland, OR). Results were presented as the percentage of positive cell populations.

### FACS-sorting of skin cells

Skin cell suspensions were resuspended in 100 μL of flow buffer per 10^7^ cells and transferred to polypropylene FACS tubes (BD Falcon, 352063). The cells were then stained with 5 μL of CD45 APC-H7 antibody (clone 2D1) per 10^7^ cells and incubated for 30 minutes at 4℃ in the dark. After staining, the cells were washed by diluting in the flow buffer and centrifuging at 500 x g for 5 minutes. The cells were resuspended in 1 mL of sort buffer per 20×10^6^ cells with 3 μM DAPI, filtered using a 100 μm cell strainer, and sorted using a BD FACSAriaII Sorter with a 100 μm fluidics nozzle. Cells isolated from FACS gates shown in Figure.S2A were then profiled by scRNA-seq.

### Single-cell RNA-sequencing

Single-cell libraries were generated using the Chromium Controller and Single Cell 5’ Library & Gel Bead Kit (10x Genomics, PN-1000006). The cell suspension was loaded onto the Chromium single cell controller (10x Genomics) to generate single-cell gel beads in the emulsion according to the manufacturer’s protocol. Up to 10,000 cells were added to each channel, and the target cell number recovered was estimated to be about 6,000 cells. Captured cells were lysed and the released RNA was barcoded through reverse transcription in individual Gel Beads-in-emulsions (GEMs).

Reverse transcription was performed on a S1000TM Touch Thermal Cycler (Bio Rad) at 53°C for 45 min, followed by 85°C for 5 min, and hold at 4°C. The cDNA was generated and then amplified, and its quality assessed using an Agilent 4200 (performed by CapitalBio Technology, Beijing). The resulting library was subsequently sequenced using the Illumina Novaseq 6000 platform in a 150-bp paired-end (PE150) configuration, with a sequencing depth of at least 100,000 reads per cell.

### scRNA-seq data processing

Preprocessing of the scRNA-seq data was performed using Cell Ranger (v5.0.1) from 10x Genomics. All transcriptome samples were aligned to the human reference genome assembly refdata-gex-GRCh38-2020-A using the Cell Ranger command “count”. The expression data were loaded into R (v4.0.3) and further analyzed using the Seurat package (v4.3.0) [13]. Cells with fewer than 200 genes, or of which the mitochondrial gene ratio was more than 5% in PBMCs or 20% in tissue samples, were regarded as abnormal and were filtered out. Gene counts were normalized using sctransform method in Seurat[14]. Dimensionality reduction was performed using PCA. To retain more than 85% cumulative variance, the Leiden algorithm was used to detect clusters in the first 23 principal components for PBMC data and separately in the first 15 principal components for skin data [15]. We used LIGER for batch-effect correction for skin data[16].

### Differential expression analysis

Detection of genes differentially expressed between the PBMC samples from psoriasis and healthy control conditions was performed using the sample-based method (the pseudobulk method of edgeR) provided by the Libra R package[17]. For skin data, a cell-based approach utilizing the Wilcoxon rank test (the default method in Seurat) was employed due to the limited number of samples per condition (only two).

### Gene Set Enrichment Analysis (GSEA)

GSEA was performed using R package clusterProfiler (version 4.0), to investigate biological functions and mechanisms that may be different in psoriasis.

### Weighted Gene Co-expression Network Analysis (WGCNA)

Filtered, scaled gene expression data of lesional samples were used as input for a biweight-midcorrelation signed network which was constructed using the hdWGCNA package (version 0.2.17) in R[18]. This gene co-expression network was constructed with the following parameters: k = 10, max_shared = 5, soft power threshold = 9, sampleForScalingFactor = 1000, deepSplit = 4, minModuleSize = 20, mergeCutHeight = 0.1, corType = “bicor”, networkType = “signed”.

### Statistical analysis

We used the Chi-square test or Fisher’s exact test for non-continuous variables and ANOVAs for continuous variables to compare the clinical characteristics. *P* values were adjusted with the Benjamini-Hochberg method for multiple hypothesis testing and adjusted *P* values <⍰0.05 were considered as significant. We computed the *Spearman* correlation between gene expression and microbial abundance. All statistical analyses were performed using the stats package (version 4.1.2) in R (version 4.0.3).

## Results

### Numbers and gene expression profiles of circulating monocytes and pDCs are affected in psoriasis

Our study utilized five datasets (Figure.1A). For the single-cell transcriptome data, we sequenced PBMCs from eleven psoriasis patients and eight healthy individuals, resulting in a total of 84,252 cells, with an average of 4,434 cells per sample (Figure.1B). We created a cells-by-genes expression matrix and performed dimensionality reduction by uniform manifold approximation and projection (UMAP) and graph-based clustering, and identified eight clusters (Figure.1C, Figure.S1A, S1B). Along with the expression of cell-type specific marker genes, we manually annotated the clusters using their respective most highly differentially expressed genes (DEGs) (Figure.S1C, S1D). To investigate the cells of different lineages with their respective optimal resolutions, we separately analyzed the myeloid cells and lymphoid cells and performed dimensionality reduction and clustering with different strategies for each of them.

Among the myeloid cells, we identified 5 clusters, for which we subsequently analyzed the disease-driven alterations in abundance (Figure.2A-2C, Figure.S1E, S1F) and gene expression profile (Fig.2D-2I). Patients with psoriasis had significantly higher proportions of CD14^+^ monocytes, plasmacytoid dendritic cells (pDCs), and hematopoietic stem and progenitor cells (HSPCs) in PBMCs than healthy controls (Figure.2B). Also the proportion of (total) monocytes and the ratio of CD14^+^ to CD16^+^ monocytes were increased in psoriasis (Figure.2C). Proportions of CD16^+^ monocytes and classical DCs (cDCs) were not significantly different between patients and controls (Figure.2B).

**Figure 2.**
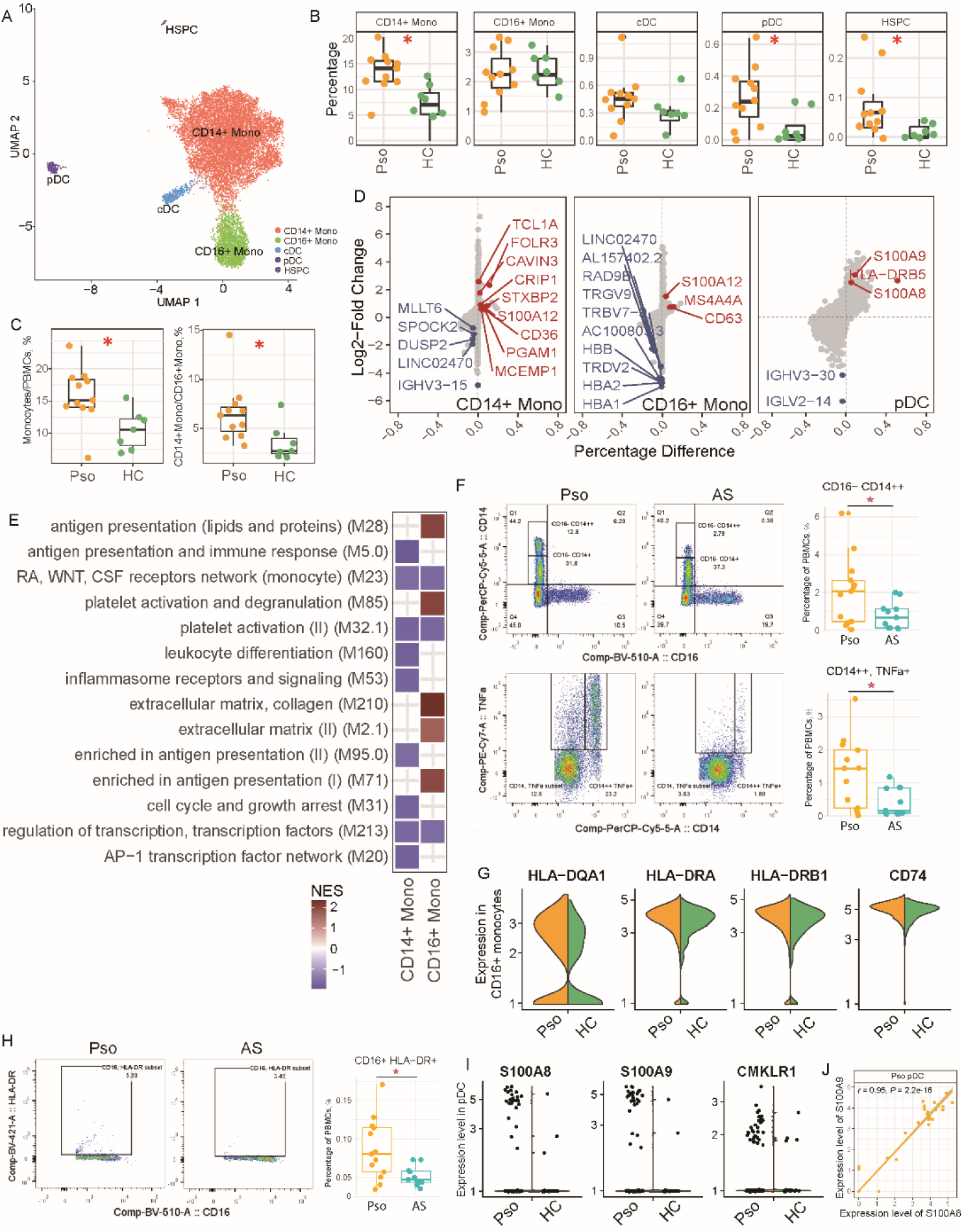
Functional annotation of circulating myeloid cells. A. UMAP visualization of myeloid cells. The pDC, and monocyte and cDC subsets were extracted from figure 1B, after which we re-ran the UMAP for these subsets only. The annotations for cell clusters are based on the DEGs of clusters (Figure S1E) and the expression of marker genes (Figure S1F). B. Abundance of myeloid cell subsets in psoriasis (Pso) and healthy controls (HC), expressed as percentage of PBMCs. Two-sided Wilcoxon test. \**P* < 0.05. C. Percentage of total monocytes (i.e. CD14^+^ monocytes plus CD16^+^ monocytes) in total PBMCs, and the ratio of CD14^+^ monocytes over CD16^+^ monocytes, in psoriasis and healthy controls. Two-sided Wilcoxon test. \**P* < 0.05. D. Condition-dependent changes in gene expression for CD14^+^ monocytes, CD16^+^ monocytes and pDCs. The top 10 upregulated and downregulated DEGs with adjusted *P* < 0.05 (LRT test with pseudobulk method) at each end are labeled with the corresponding gene names. The genes in red are upregulated DEGs in psoriasis, while those in blue are downregulated. E. GSEA pathway analysis of the differentially expressed genes in psoriasis across myeloid cell subsets, compared to healthy controls. F. The flow cytometric scatter plots of CD14^high^ monocytes (upper panel) and CD14^high^ TNF^+^ monocytes (lower panel) in psoriasis and ankylosing spondylitis. And the percentage of CD14^high^ monocytes (upper panel) and the percentage CD14^high^ TNF^+^ monocytes (lower panel) in total PBMCs in psoriasis and ankylosing spondylitis. Two-sided Wilcoxon test. \**P* < 0.05. G. Expression profiles of marker genes of MHC class II in CD16^+^ monocytes. These genes were differentially expressed between the psoriasis and healthy conditions (adjusted *P* value < 0.05). H. The flow cytometric scatter plots of CD16^+^ HLA-DR^+^ monocytes in psoriasis and ankylosing spondylitis. Percentage of CD16^+^ HLA-DR^+^ monocytes in total PBMCs. Two-sided Wilcoxon test. *P < 0.05. I. Expression profiles of S100A8, S100A9 and CMKLR1(ChemR23) in pDCs. These genes were differentially expressed between the psoriasis and healthy conditions (adjusted *P* value < 0.05). J. Co-expression of S100A8 and S100A9 in circulating pDCs in psoriasis.

Comparing the expression levels of individual genes between conditions, we found that the expression of several S100 family genes was upregulated in circulating myeloid cells from psoriasis patients. For instance, S100A12 was upregulated in both CD14^+^ and CD16^+^ monocytes (Figure.2A). Using GSEA, we found that especially genes related to “*inflammasome receptors and signaling”* and “*antigen presentation and immune response”*were suppressed in CD14^+^ monocytes in psoriasis (Figure.2E, Figure.S1I), while genes related to antigen presentation of CD16^+^ monocytes were enriched in psoriasis (Figure.2E, Figure.S1J). We performed flow cytometry analysis on PBMCs from patients with psoriasis and ankylosing spondylitis (AS) and identified two distinct populations of CD14^+^ monocytes: CD14^low^ and CD14^high^. The CD14^high^ monocytes were more prevalent in psoriasis patients compared to those with AS. Additionally, TNF expression in the CD14^high^ monocytes was elevated in psoriasis compared to AS (Figure 2F Figure.S2). The enhanced antigen presentation by CD16^+^ monocytes in psoriasis was indicated by the significantly increased expression of MHC class II genes (HLA-DRA1, HLA-DRB1, HLA-DQA1, and CD74) (Figure 2I). And the expression of HLA-DR on CD16+ monocytes was consistently higher in psoriasis.

Besides, pDCs showed upregulated expression of S100A8 and S100A9 in psoriasis (Figure.2I). The expression of S100A8 and S100A9 was positively correlated (Figure.2J). Expression of the skin-homing marker CMKLR1 was also up-regulated in circulating pDCs of psoriasis patients (Figure.2I).

In summary, our study demonstrated an increased frequency of circulating CD14^high^ monocytes in psoriasis, along with upregulated TNF expression. Additionally, CD16+ monocytes in psoriasis patients showed enhanced antigen presentation through elevated MHC class II expression. pDCs in psoriasis also exhibited potential skin-homing and pro-inflammatory characteristics.

### Circulating memory T cells and NK cells in psoriasis show signs of a less activated phenotype

To analyze possible changes in lymphoid cells in psoriasis, we re-clustered the circulating T cell and natural killer (NK) cell subsets from PBMCs as identified in Figure.1B. This yielded 14 clusters based on the expression of marker genes and DEGs (Figure.3A, Figure.S3A, S3B). Surprisingly, the proportion of CD8^+^ T_EM_ cells in circulation dropped significantly in psoriasis (Figure.3B). Transcriptionally, none of the conventional proinflammatory genes was upregulated in memory T cells in psoriasis (Figure.3C). We extended the investigation for circulating T cells by analyzing their gene expression related to activation, differentiation, and migration patterns. Remarkably, gene expression of many T cell pathways, especially T cell activation, was lower in circulating T cells in psoriasis compared to healthy controls (Figure.3D). We then evaluated the migration potential of circulating T_EM_ cells in psoriasis by measuring the expression of skin-homing markers [26]. We found that SELPLG (also known as CLA), ITGAE (which encodes CD103), and CXCR3 were up-regulated in CD8^+^ T_EM_ cells in psoriasis (Figure.3E). To further validate these findings, we examined the flow cytometry profile and found that while the abundance of activated CD8+ T cells (CCR4+ CD8+ T cells) was lower in psoriasis (Figure.3F, Figure.S4), the expression of CLA on these CCR4+ CD8+ T cells was higher(Figure.3G, Figure.S4). This collective pattern suggested an enhanced propensity of circulating T_EM_ cells in psoriasis to migrate towards the skin.

**Figure 3.**
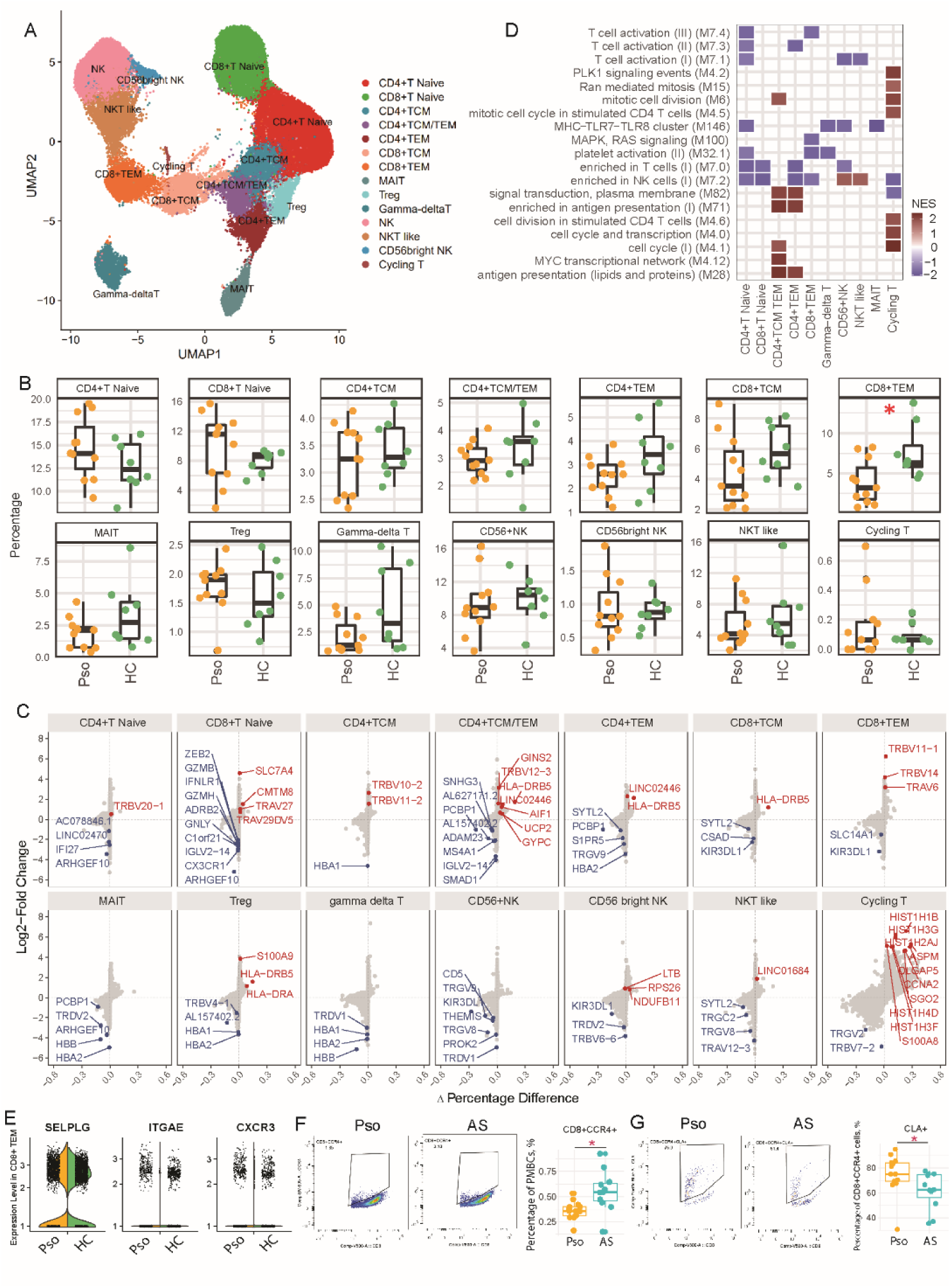
Analysis of circulating lymphoid cells. A. UMAP visualization of T and NK cells. The data on T and NK cell subsets were extracted from figure 1A and re-run in a new UMAP analysis. The annotations for cell clusters are based on the DEGs of clusters (Figure.S3A) and the expression of marker genes (Figure.S3B). B. Abundance of T and NK cell subsets in psoriasis and healthy controls, expressed as percentage of PBMCs. Two-sided Wilcoxon test. * *P* < 0.05. C. Condition-dependent changes in gene expression for each cell type. The top 10 upregulated and downregulated DEGs with adjusted *P* < 0.05 (LRT test with pseudobulk method) at each end are labeled with the corresponding gene names. The genes in red are upregulated DEGs in psoriasis, while those in blue are downregulated. D. GSEA pathway analysis of the differentially expressed genes of psoriasis across T and NK cell subsets, compared to healthy controls. E. The expression of skin-homing maker genes SELPLG(CLA), ITGAE(CD103) and CXCR3 in CD8^+^ T cells between conditions. F. The flow cytometric scatter plots of CD8^+^ CCR4^+^ T cells in psoriasis and ankylosing spondylitis. Percentage of CD8^+^ CCR4^+^ T cells in total PBMCs. Two-sided Wilcoxon test. *P < 0.05. G. The flow cytometric scatter plots of CD8^+^ CCR4^+^ CLA^+^ T cells in psoriasis and ankylosing spondylitis. Percentage of CLA^+^ cells in CD8^+^ CCR4^+^ T cells. Two-sided Wilcoxon test. *P < 0.05.

For NK cells, we identified three subsets in the circulation: classical NK cells, CD56 bright NK cells, and NKT like cells (Figure.3A). No significant differences were found in the percentages of these subsets between psoriasis patients and healthy individuals (Figure.3B). Nevertheless, we found some differences in their gene expression levels. Similar to the DEG profile of memory T cells, we did not observe any significant upregulation of genes of interest in psoriasis. Instead, we observed downregulation of KIR3DL1 in classical NK cells and CD56 bright NK cells (Figure.3C), and downregulation of some genes related to pro-inflammatory pathways, such as the MHC-TLR7-TLR8 cluster, in classical NK cells (Figure.3D).

We next investigated the profile of B cells in the circulation. Re-clustering of the B cell gene expression data yielded four clusters (Figure.S5A-S5C). Although the proportions of the four B cell subsets (i.e. naive, memory, intermediate and plasma B cells) did not differ significantly between individuals with and without psoriasis, the expression of genes associated with immune responses of these B cell subsets was suppressed in psoriasis (Figure.S5D-S5F).

In search of further support for the observed differences in gene expression in PBMCs between participants with and without psoriasis, we conducted the same GSEA on two public bulk RNA-seq datasets of PBMCs from psoriasis patients. The validation results confirmed our findings: In PBMCs from psoriasis patients, there was higher expression of marker genes associated with monocytes, fewer genes activated in T cells, and similar gene expression profiles in B cells compared to healthy controls(Figure.S6A∼S6D).

### Proinflammatory gene expression is upregulated in most myeloid subsets in psoriatic lesions

Of the eleven psoriasis patients in this study, two donated psoriatic lesional and non-lesional tissue samples, which were used to study the impact of psoriasis on the skin at single-cell resolution. With skin single-cell transcriptome analysis, after prefiltering, we obtained the lymphoid and myeloid clusters from CD45^+^ cell suspensions (Figure.S8A-S8D). The cell counts of both myeloid and lymphoid cells were increased in lesions, but the ratio of myeloid to lymphoid cells tended to be lower in lesions than in non-lesions (Figure.S8E, S8F). Similar to the data processing of PBMCs, we subsequently analyzed the gene expression data of myeloid cells and lymphoid cells separately.

The myeloid cells in skin were re-clustered and re-annotated into 5 myeloid subsets (Figure.4A, Figure.S8G, S8H). Unexpectedly, the two macrophage clusters did not adhere to the M1/M2 dichotomy [27,28]. Instead, they differentiated between monocyte-derived (MONO Mac) and resident (RESINT Mac) macrophages. Monocyte-derived macrophages (expressing the monocyte markers CD14, CD68 and FCGR3A) included both M1 and M2 macrophages, based on their gene expression signatures (Figure.S8H, S8I). Resident macrophages exclusively expressed the tissue-resident markers HES1 and LYVE1[29] (Figure.S8H). Only a small population of resident macrophages exhibited an M1 or M2 signature (Figure.S8I). The ratio of monocyte-derived to resident macrophages was higher in lesions than in non-lesions (Figure.4B).

**Figure 4.**
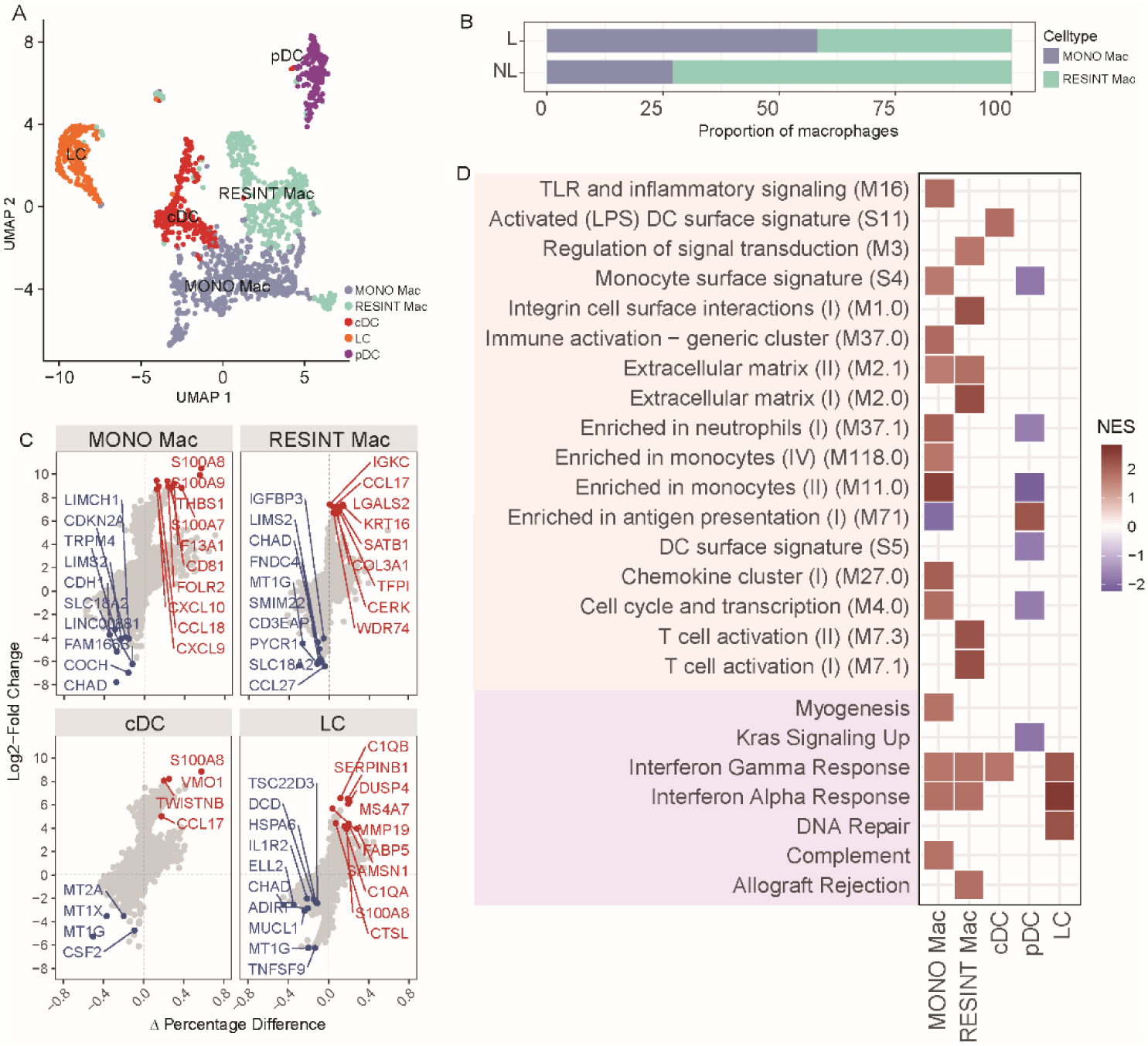
Analysis of myeloid cells in lesional and non-lesional skin. A. UMAP visualization of myeloid cells in skin. The data on myeloid cells were extracted from Figure.S8C and re-run in a new UMAP analysis. The annotations for cell clusters are based on the expression of marker genes (Figure.S8G) and the DEGs of clusters (Figure.S8H). B. Bar charts showing the normalized cell counts of myeloid cell subsets in lesions (L) and non-lesions (NL). C. Condition-dependent changes in gene expression for each cell type. The top 10 upregulated and downregulated DEGs with adjusted P < 0.05 (Wilcoxon rank test) at each end are labeled with the corresponding gene names. The genes in red are upregulated DEGs in lesions, while those in blue are downregulated. D. GSEA pathway analysis of the DEGs of myeloid cell subsets in lesions compared to non-lesions. The upper panel was annotated with BTM and the lower panel with GO.

At the transcript level, similar to what we observed for peripheral myeloid cells, the expression of S100A7/8/9 was upregulated in most of the myeloid subsets in lesions. Monocyte-derived macrophages in lesions strongly expressed the genes for the IFN-γ-inducible ligands CXCL9 and CXCL10 (Figure.4C, S8L). Consistent with the DEG profile, the expression of most genes related to immune responses was enhanced in myeloid subsets in lesions, based on both BTM and GO (Gene Ontology) annotation (Figure.4E). For instance, gene expression of the IFN-α and IFN-γ pathways were both elevated in most of the myeloid cell subsets in psoriasis.

Taken together, we observed that psoriatic lesions exhibit an increased presence of monocyte-derived macrophages. The gene expression of most myeloid subsets in lesions displayed a more pro-inflammatory profile compared to those in non-lesions.

### Lymphoid subsets are broadly increased in numbers and activated in psoriatic lesions

In skin, we identified ten subtypes of lymphoid cells: activated CD8^+^ T_RM_, resting CD8^+^ T_RM_, CD8^+^ T_EM_, CD4^+^ T_RM_, CD4^+^ T_EM_, Treg, MAIT, innate lymphoid cells (ILCs), NK, and cycling T cells. The remaining cells, including a few CD4^+^ T_EM_ Cells, B cells and some other cells were clustered as an undefined subset (Figure.5A, Figure.S9A, S9B). Similar to what we observed for skin myeloid cells, the cell counts of all these lymphoid subsets tended to be higher in lesions (Figure.S9C, S9D). At the transcript level, the expression of proinflammatory cytokines and chemokines by memory T cell subsets was also significantly higher in lesional than in non-lesional skin samples (Figure.5B). Also the expression of IL17A and IL22, which are associated with psoriasis[2], was increased in most cell subsets from lesional skin. Th17 and Tc17 cells were mainly T_RM_ cells, and not T_EM_ cells (Figure.5C). Due to the limited number of IL17-producing cells that were identified, and their heterogeneous nature within clusters, we chose not to label any cluster as either Th17 or Tc17 (Figure.S9C), unlike other single-cell studies of psoriasis [8,9]. At the cellular pathway level, most of the genes for the pathways related to immune responses were upregulated in psoriatic lesions. For instance, the genes for “*T cell activation*, *MHC-TLR7-TLR8 cluster”* and “*interferon alpha* /*gamma pathways”* were upregulated across most of the memory T-cell subsets in lesional skin samples (Figure.5D).

**Figure 5.**
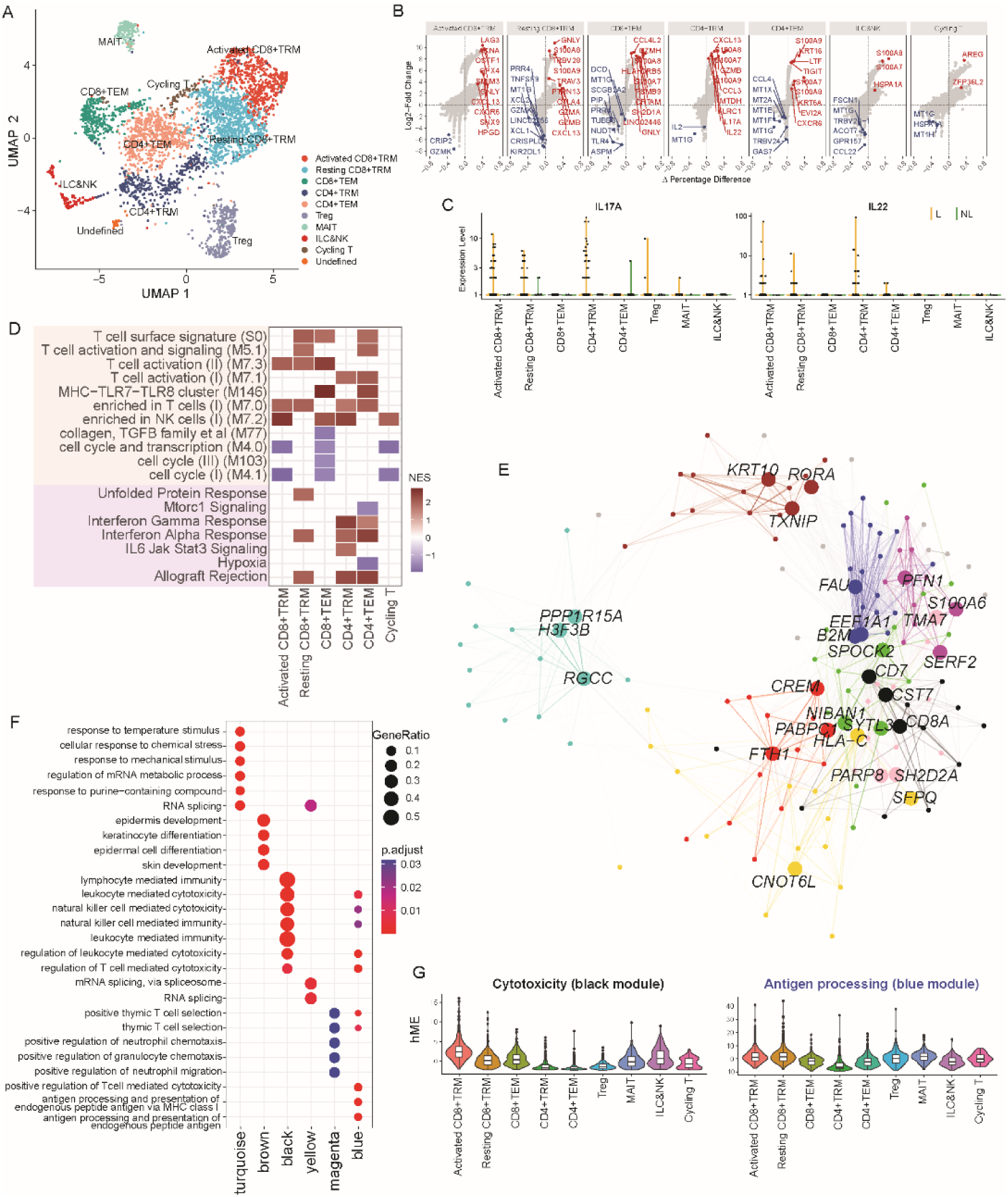
Analysis of lymphoid cells in lesional and non-lesional skin. A. UMAP visualization of skin lymphoid cells. The data of skin lymphoid cell subsets were extracted from Figure.S8C and re-run in a new UMAP analysis. The annotations for cell clusters are based on the DEGs of clusters (Figure.S9A) and on the expression of marker genes (Figure.S9B). B. Condition-dependent changes in gene expression for each cell type. The top 10 upregulated and downregulated DEGs with adjusted *P* < 0.05 (Wilcoxon rank test) at each end are labeled with the corresponding gene names. The genes in red are upregulated DEGs in lesional skin, while those in blue are downregulated. C. Expression of the psoriasis-related proinflammatory maker genes IL17A and IL22 across skin lymphoid cell subsets in lesional and non-lesional skin. D. GSEA pathway analysis of the differentially expressed genes across different lymphoid cell subsets in lesional compared to non-lesional skin. The upper panel was annotated with BTM and the lower panel with GO. E. WGCNA network of the top 15 hub genes of each module, where each node represents a gene and each edge a co-expression relationship. The top 3 hub genes of each module are labeled with their gene names. The opacity of edges was scaled by the strength of the co-expression relationship. F. GO annotation of the genes in each module. G. Harmonized module eigengenes (hMEs) of the black and blue modules across different lymphoid cell types in the skin.

Since the expression of most proinflammatory genes differed between psoriasis patients and healthy individuals, we were able to find clusters of highly correlated genes (modules) with certain biological functions that are overrepresented in psoriasis, and to identify the most important genes (hub genes) within each module. Using Weighted Gene Co-expression Network Analysis (WGCNA), we constructed gene co-expression networks across lymphoid cells from psoriatic lesions and identified nine modules (Figure.5E, Figure.S9D, S9E). Two of these are considered to have specific biological functions with GO annotation related to the immune responses: the cytotoxicity module (black) with hub genes CD8A, CST7 and CD7; and the antigen processing module (blue) with hub genes B2M, EEF1A1 and FAU (Figure.5E, 5F). To compare the expression level of these two modules across the different skin lymphoid cell subsets, we measured the eigengenes, which can be interpreted as the representative expression profile of each module. Activated CD8^+^ T_RM_ cells showed strong expression of both signatures. Resting CD8^+^ T_RM_ cells showed similar gene expression profiles related to antigen processing as activated CD8^+^ T_RM_ cells, but with lower expression levels of cytotoxicity-related genes (Figure.5G). In addition to T cells, ILCs and NK cells also expressed higher levels of S100 family genes in lesions (Fig 5B). ILCs and NK cells showed comparable activity in the cytotoxic module compared to CD8+ T cells, but they exhibited much weaker activity in the antigen processing module(Fig 5G).

In summary, we found that the gene expression profiles of skin lymphoid subsets are indicative of activation in psoriatic lesions. Our results demonstrate the gene expression pattern associated with cytotoxicity and antigen processing in CD8^+^ T_RM._ Lastly, ILCs and NK cells were present in the inflammatory signature observed in psoriatic lesions.

## Discussion

Understanding the aberrations of immune cells in psoriasis is important for identifying potential new targets for treatment that can help restore normal immune function and alleviate symptoms. To address this, we utilized single-cell transcriptome analysis to study the phenotypes and gene expression alterations of multiple immune cell subsets in the blood and skin in psoriatic and non-psoriatic conditions. We found clear alterations in the gene expression patterns of both myeloid and lymphoid cells in psoriasis, which were partially overlapping and partially unique in blood and skin.

In comparison to healthy controls, we found that patients with psoriasis had higher proportions of CD14^+^ monocytes, which themselves had a reduced pro-inflammatory profile. For instance, the expression of TLR4 was lower in CD14^+^ monocytes from patients with psoriasis compared to healthy controls. For CD16^+^ monocytes in psoriasis, we observed an increased expression of genes related to antigen presentation. The increased expression of MHC class II and CD74 observed for CD16^+^ monocytes in psoriasis suggests a potential role of these cells in presenting antigens to T cells[34].

pDCs are known for their substantial secretion of cytokines, including interferons with antiviral and immunomodulatory properties. Several studies have documented the gathering of pDCs within psoriatic lesions, where they actively participate in local immune reactions [35,36]. In the circulation, we observed an elevated prevalence of pDCs in psoriasis patients. These pDCs exhibited higher expression of genes related to skin homing (CMKLR1) and pro-inflammatory responses (S100A8 and S100A9), and the expression of S100A8 and S100A9 was positively correlated. The positive correlation between the expression of S100A8 and S100A9 suggests that they may assemble into S100A8/S100A9 dimers (calprotectin) [37]. In previous studies of chronic inflammatory diseases, calprotectin have been observed [38,39]. Calprotectin can bind to TLR4 and enhance antiviral activity in target cells [40]. The positive correlation between the expression of S100A8 and S100A9 that we observed thus opens up the possibility for them to form calprotectin. These attributes of circulating pDCs in psoriasis suggest that they may be more prone to relocate to the skin and engage in localized inflammation.

Also the profiles of circulating T cells were altered in psoriasis, with reduced expression of genes related to cytotoxicity and activation. It is well-established that the frequency and activation levels of specific T cell subsets differ between individuals with and without psoriasis [2]. We discovered that circulating memory T cells in psoriasis patients were reduced in numbers and exhibited a less functional phenotype at the transcript level. This signature was confirmed by our reanalysis of two publicly available bulk RNA-seq datasets. We propose that this T-cell “suppression” profile in psoriasis may be caused by migration of memory T cells from the circulation to the lesions. Recent studies have reported that the proportions of immune cells in the circulation are affected by age [41,42]. To rule out a potential bias of age in our study, we also examined the impact of age on the abundance of all cell subsets in PBMCs. No significant correlation was observed after adjusting for multiple testing (Figure.S5).

In our cohort, we were fortunate to have two patients who were willing to donate lesional and non-lesional skin samples. In line with a previous study[11], we found upregulation of most myeloid cells, both in terms of their quantity and their gene signature related to immune responses, when comparing lesional to non-lesional skin. In contrast to the study of Zhang et al. [10], our findings did not show any clusters of M1 or M2 macrophages. Instead, we identified two distinct clusters of macrophages based on their origin: monocyte-derived macrophages and tissue-resident macrophages. The number of monocyte-derived macrophages was higher in psoriatic lesions.

The T cell profile in skin showed a hyperinflammatory state at the transcript level in psoriatic lesions: T cells were both increased in numbers and more activated in lesions compared to non-lesions. Some recent single-cell studies focused on Th17 and Tc17 cells, which are the canonical cell subsets contributing to psoriasis [8,9]. In line with these studies, we found an enrichment of IL17-producing cells in lesions. These IL17-producing cells showed a strong tissue-resident memory T cell signature, suggesting that these cells have migrated to and taken up long-term residence in the tissues affected by psoriasis. We noted that resting CD8^+^ T_RM_ cells in lesions may retain certain immune functions at transcript level, especially related to antigen processing.

Our data will hopefully serve as a useful reference for future studies on the links between immune cells in the circulation and skin in psoriasis. In addition to deciphering the role of the innate and adaptive immune system in the pathophysiology of psoriasis, future studies should examine the role of migration of monocytes and pDCs from the circulation to the skin. A limitation of our study is the small number of patients included in the investigation of paired skin and blood samples. Furthermore, since it can be challenging to measure transcript levels of canonical markers (for instance, cytokines) that are often used to define cell types based on scRNA-seq data, additional methods are necessary to enhance our comprehension of single-cell observations. Finally, our observations are limited to the transcriptional level. The functional implications of these observations require further investigation.

In conclusion, our single-cell study delineates the immune landscape in psoriasis. Our findings suggest that psoriasis involves multiple molecules and pathways, which may affect different aspects of the immune response, including antigen presentation, cell activation and expansion, and cytokine secretion and signaling. This also poses a challenge for developing effective treatments for psoriasis, as targeting one molecule or pathway may not be enough to control the disease.

## Acknowledgements

We would like to thank all the patients and healthy donors for participating in the study. We extend our gratitude to Dr. Emmerik Leijten (LangeLand Hospital) for his valuable input and feedback on the manuscript, as well as his prior contributions to the flow cytometry data used in this study. Our sincere thanks also go to Prof. Carl Goodyear (University of Glasgow) for his valuable input on an earlier version of the manuscript.

## Author Contributions

Conceptualization: CL, JD

Methodology: BG, AP, JB, JD

Data curation: JD, SY

Formal analysis: JD

Investigation: WY, JD

Funding acquisition: CL

Project administration: JB, AP, BG, CL, TR

Supervision: JB, AP, BG

Resources: CL, MON, SY, JD, JY,

Writing – original draft: JD

Writing – review & editing: JB, BG, AP, DB, MON, JD

## Data And Materials Availability

The single-cell transcriptome data used in the analysis is available in GSA database (HRA010144).

## Supplementary materials

**Supplementary Table 1.** Characteristics of the participants in scRNA-seq analysis.

|  | **Pso (n = 11)** | **HC (n = 8)** | **P value** |
| --- | --- | --- | --- |
| **Sex** |  |  |  |
| Female | 1 (9.1%) | 2 (25%) | 0.55 |
| Male | 10 (90.9%) | 6 (75%) |  |
| **Age** |  |  |  |
| Median [Min, Max] | 35 [21, 57] | 38 [22, 51] | 0.73 |
| Median [Min, Max] are presented for continuous values. Frequencies (proportion) are presented for categorical values. We used the Fisher's exact test for comparison of sex, and Wilcoxon test for comparison of age. |  |  |  |

**Supplementary Table 2.** Characteristics of the participants in flow cytometry analysis.

|  | **Pso (n = 13)** | **AS (n = 11)** | **P value** |
| --- | --- | --- | --- |
| **Sex** |  |  |  |
| Female | 7 (53.8%) | 3 (27.3%) | 0.359 |
| Male | 6 (46.2%) | 8 (72.7%) |  |
| **Age** |  |  |  |
| Median [Min, Max] | 41 [20, 63] | 47 [26, 54] | 0.706 |
| Median [Min, Max] are presented for continuous values. Frequencies (proportion) are presented for categorical values. We used the Chi-Squared test for comparison of sex, and Wilcoxon test for comparison of age. |  |  |  |

**Supplementary Figure 1.**
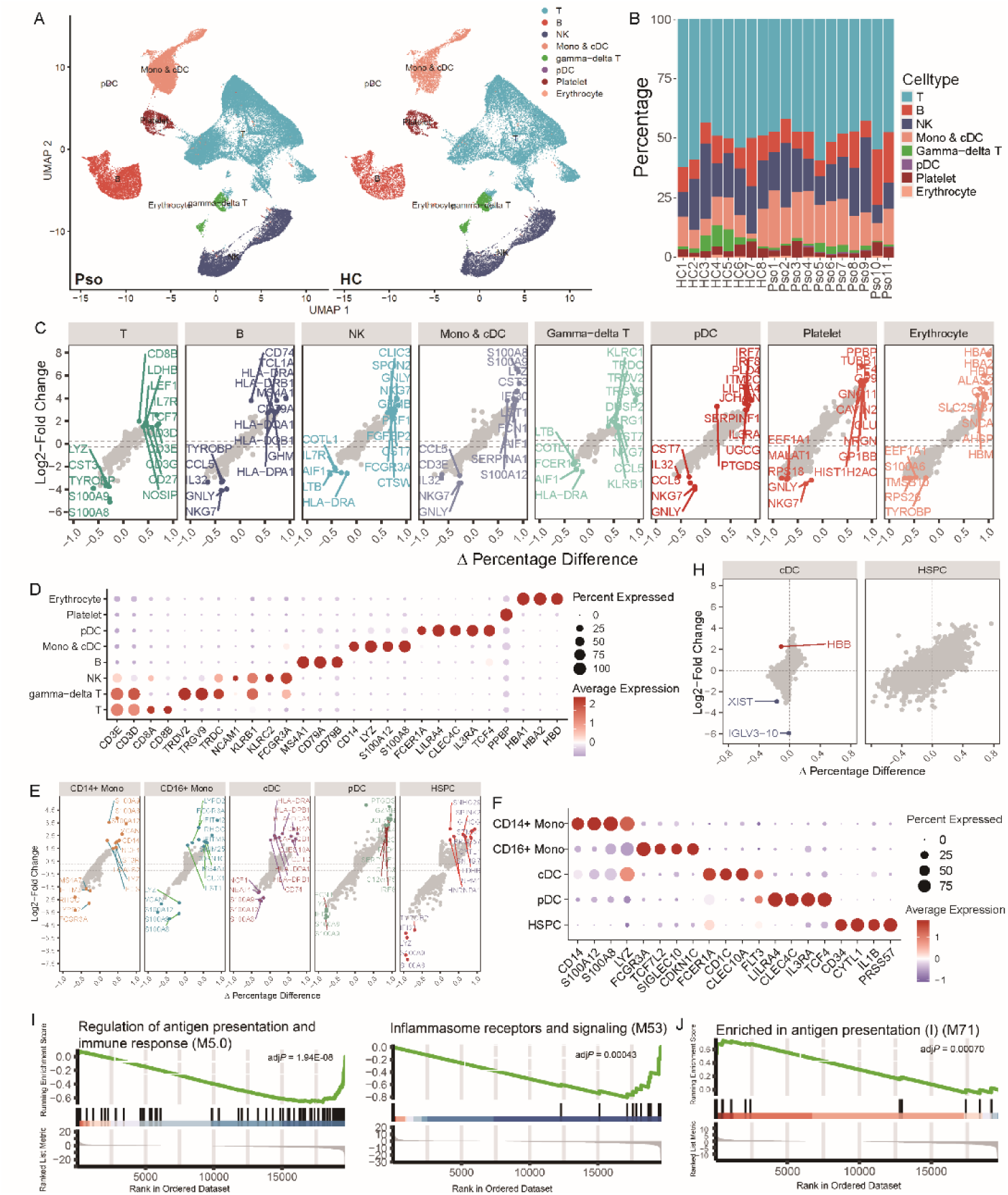
Analysis of circulating myeloid cells. A. UMAP visualization of PBMCs from psoriasis patients (left) and healthy controls (right). B. Proportions of primary cell subsets in all individuals. C. Expression of signature genes of the primary cell subsets. Log2fold changes between cell subset of interest and all other cells. Percentage difference = (proportion of cells expressing the gene of interest in the cell subset of interest) – (proportion of cells expressing the gene of interest in all other cells). D. Dot plot showing marker gene expressions across the primary cell subsets. Dot size indicates fraction of expressing cells, colored according to z-score normalized expression levels. E. Expression of signature genes of the circulating myeloid subsets. Log2fold changes between cell subset of interest and all other cells. Percentage difference = (proportion of cells expressing the gene of interest in the cell subset of interest) – (proportion of cells expressing the gene of interest in all other cells). F. Dot plot showing expression of marker genes across the circulating myeloid subsets. Dot size indicates fraction of expressing cells, colored according to z-score normalized expression levels. G. Percentage of monocytes in total PBMCs, percentage of CD16^+^ monocytes in total monocytes and the ratio of CD14^+^ monocytes to CD16^+^ monocytes between conditions. Two-sided Wilcoxon test: \**P*<0.05. H. Condition-dependent changes in gene expression for cDCs and HSPCs. The DEGs with adjusted *P* < 0.05 (LRT test with pseudobulk method) at each end are labeled with the gene names. The genes in red are significantly upregulated in psoriasis, while in those in blue are significantly downregulated. I. Visualization of functional enrichment for CD14^+^ monocytes. J. Visualization of functional enrichment in antigen presentation for CD16^+^ monocytes.

**Supplementary Figure 2.**
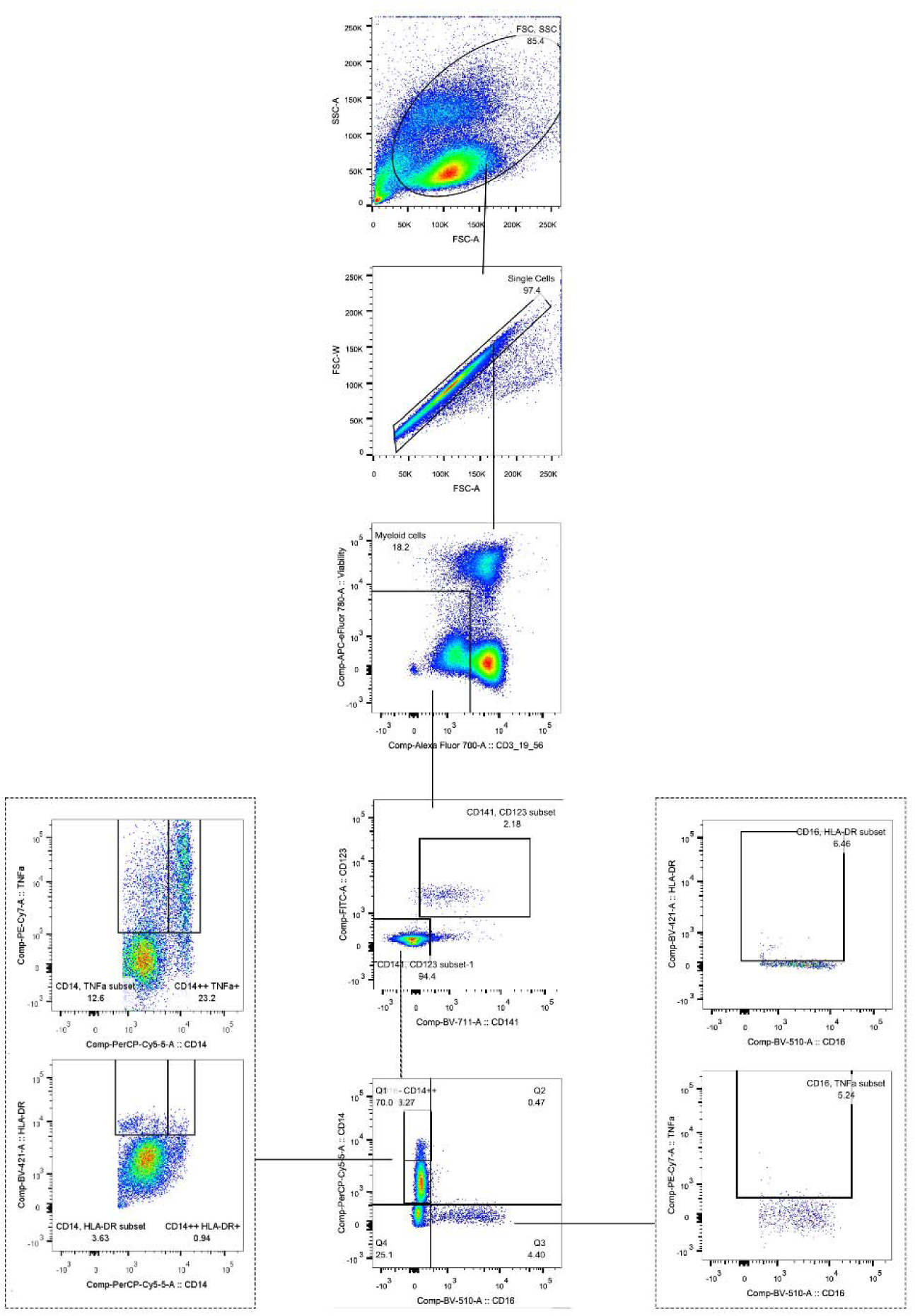
Gating strategy for identifying subsets and functions of monocytes in PBMCs.

**Supplementary Figure 3.**
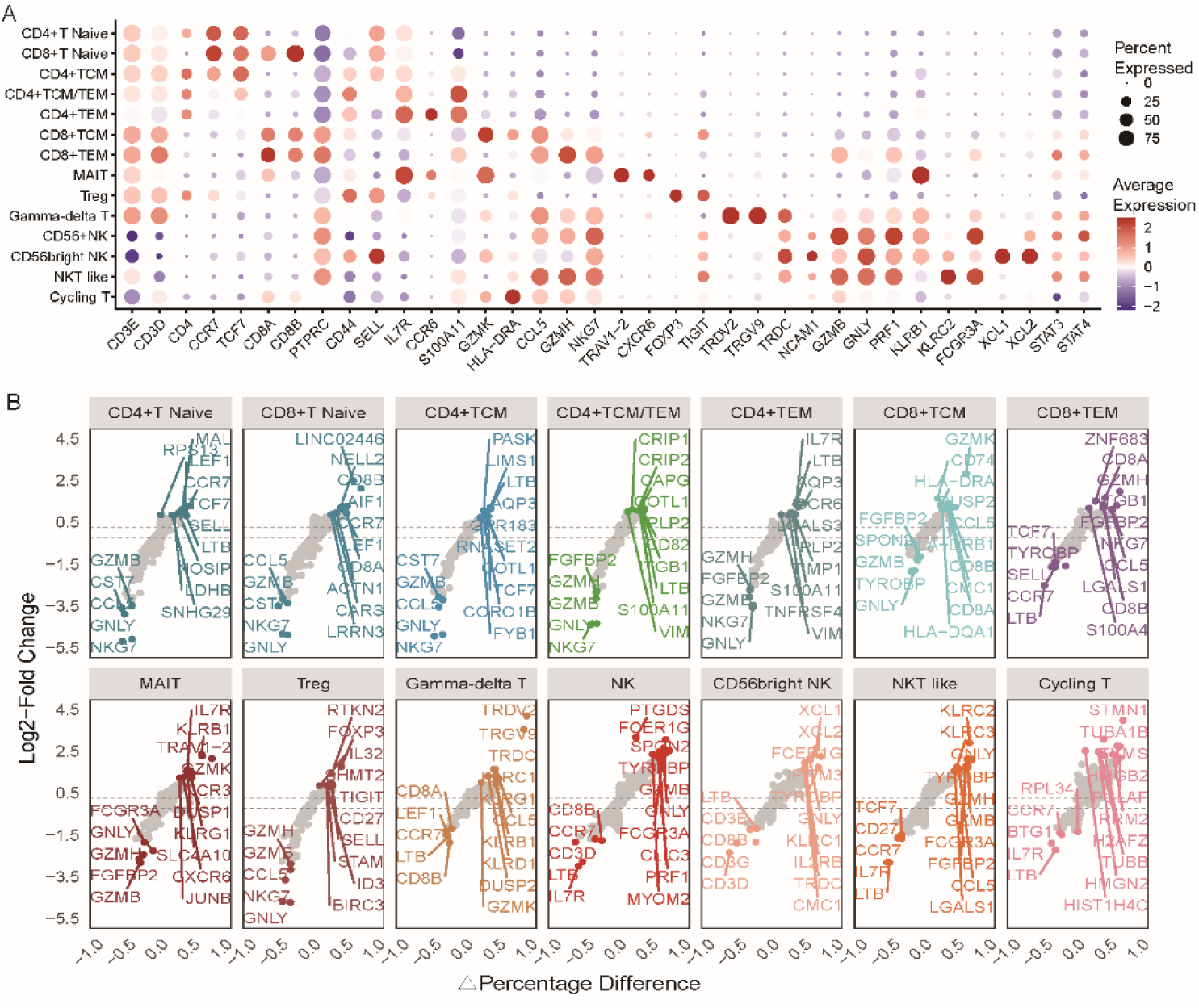
Features of circulating T and NK cell subsets. A. Dot plot showing expression of marker genes of T and NK cell subsets. Dot size indicates fraction of expressing cells, colored according to z-score normalized expression levels. B. Expression of signature genes of the T and NK cell subsets. Log2fold changes between cell subset of interest and all other cells. Percentage difference = (proportion of cells expressing the gene of interest in the cell subset of interest) – (proportion of cells expressing the gene of interest in all other cells).

**Supplementary Figure 4.**
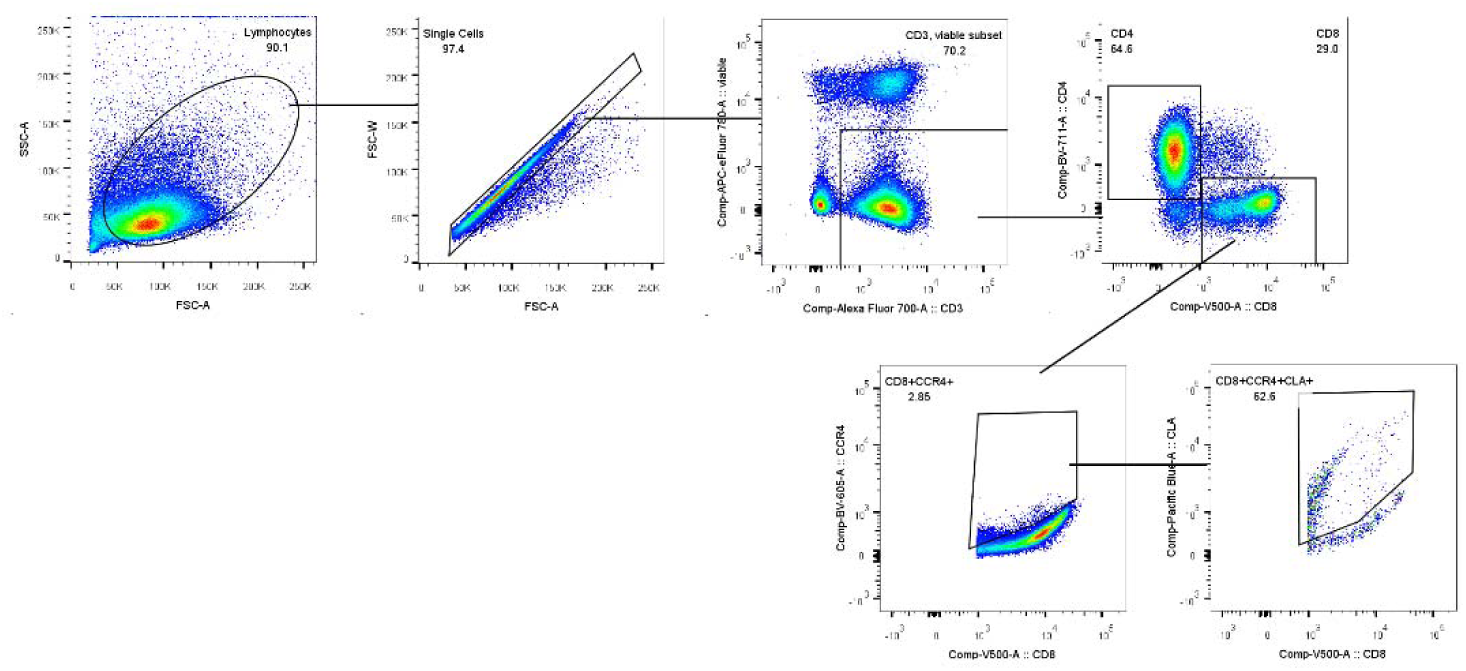
Gating strategy for identifying subsets and functions of CD8^+^ cells in PBMCs.

**Supplementary Figure 5.**
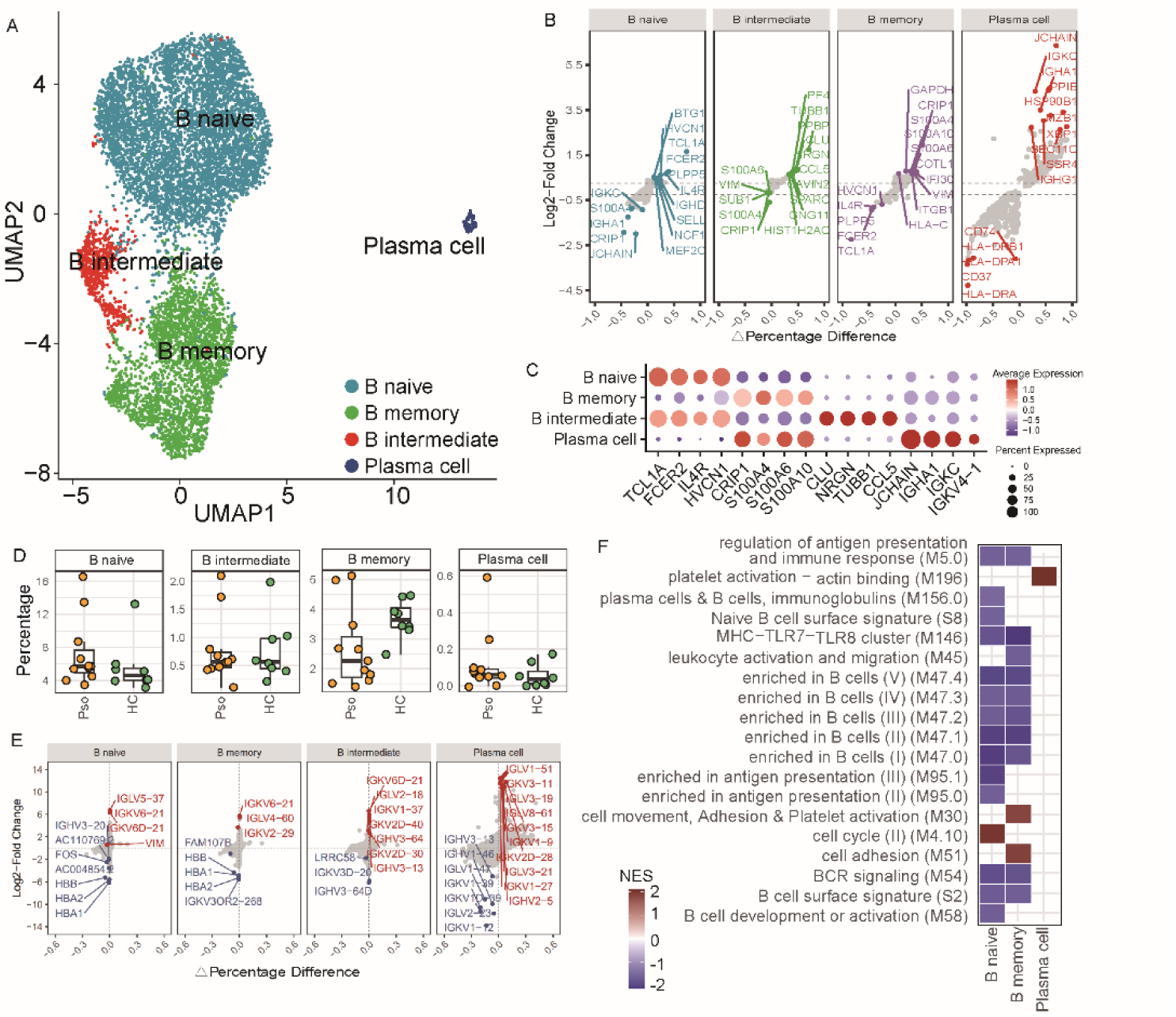
Analysis of circulating B cells. A. UMAP visualization of B cell subsets. B cells were extracted from Figure 1B and we subsequently re-ran the UMAP for this plot. The annotations for cell clusters are based on the DEGs of clusters (Figure.S3B) and on the expression of marker genes (Figure.S3C). B. Expression of signature genes of the B cell subsets. Log2fold changes between cell subset of interest and all other cells. Percentage difference = (proportion of cells expressing the gene of interest in the cell subset of interest) – (proportion of cells expressing the gene of interest in all other cells).C. Dot plot showing expression of marker genes of B cell subsets. Dot size indicates fraction of expressing cells, colored according to z-score normalized expression levels. D. Abundance of T and NK cell subsets in psoriasis patients and healthy controls. Two-sided Wilcoxon test: \**P* <0.05. E. Condition-dependent changes in gene expression for each cell type. The top 10 upregulated and downregulated DEGs with adjusted *P* < 0.05 (LRT test with pseudobulk method) are labeled with the gene names. The genes in red are upregulated in psoriasis, while those in blue are downregulated. F. GSEA pathway analysis of the differentially expressed genes of psoriasis across T and NK cell subsets, compared to healthy controls.

**Supplementary Figure 6.**
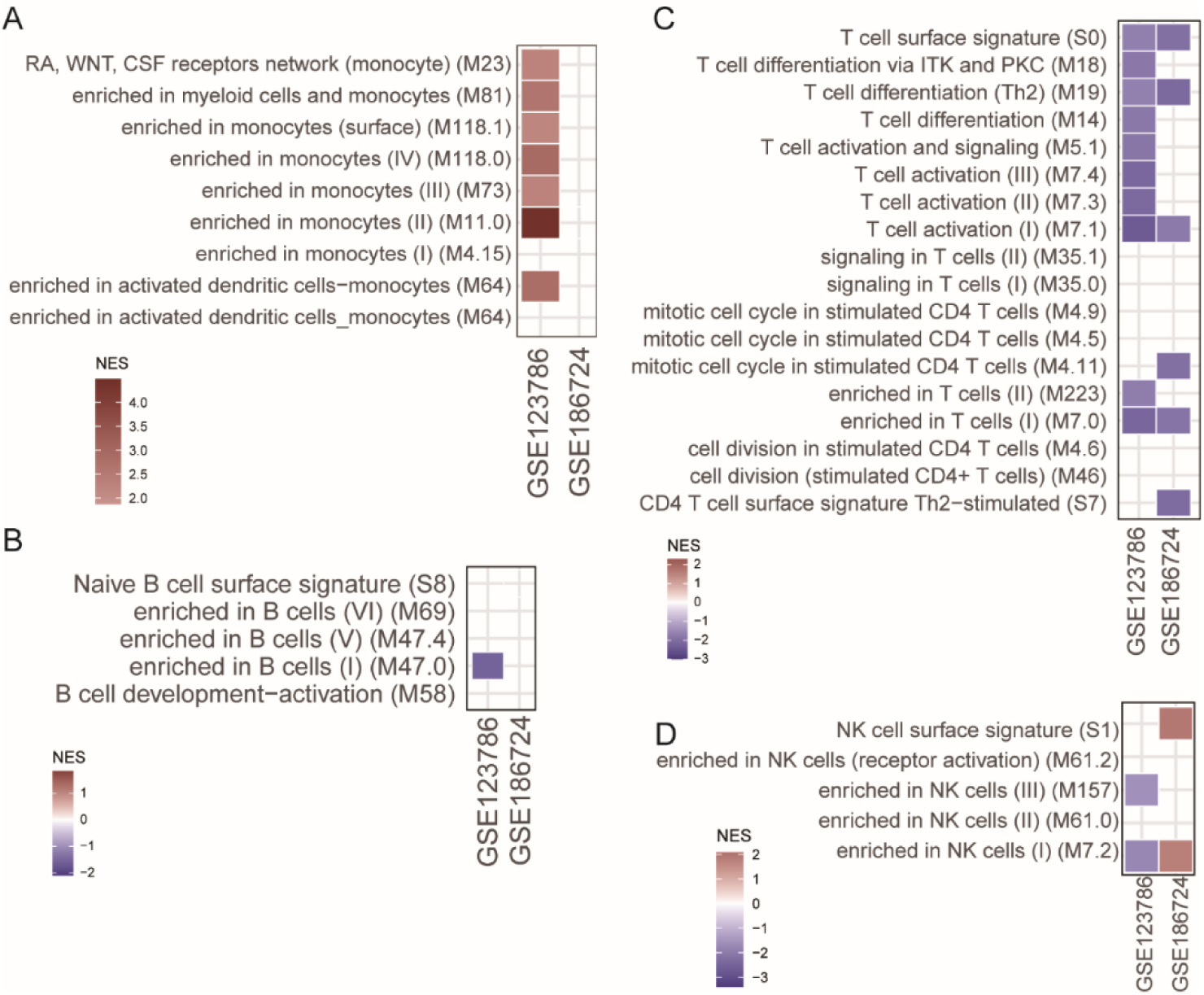
Functional enrichment of two publicly available PBMC RNA-seq datasets of patients with psoriasis (GSE123786 and GSE186724). A. Functional enrichment with BTM terms related to monocytes. B. Functional enrichment with BTM terms related to B cells. C. Functional enrichment with BTM terms related to T cells. D. Functional enrichment with BTM terms related to NK cells.

**Supplementary Figure 7.**
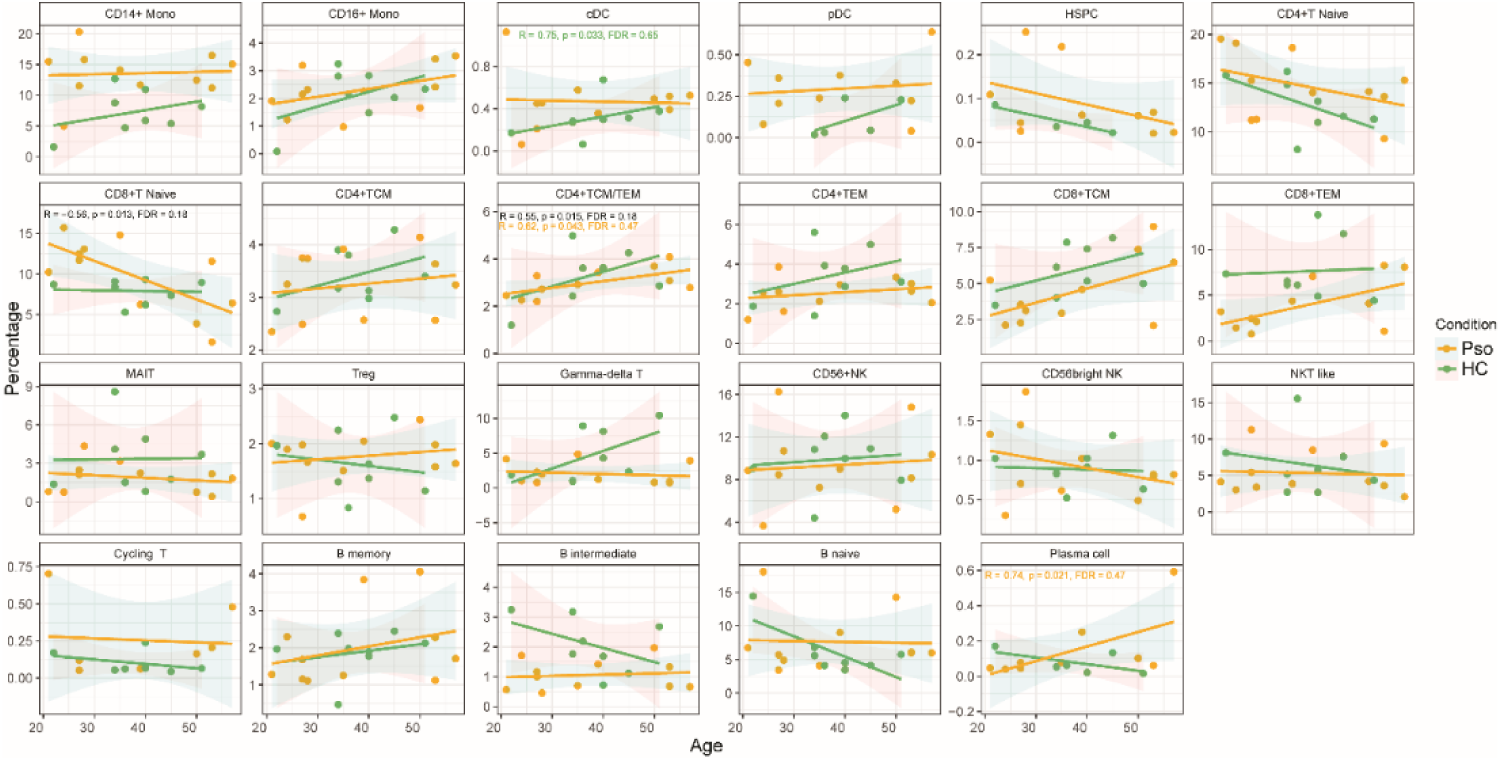
Correlation between age and proportions of circulating cell subsets for all donors. Significance was tested using the Spearman correlation test; only significant correlations are labeled with statistic data. Yellow denotes patients with psoriasis. Green denotes healthy controls.

**Supplementary Figure 8.**
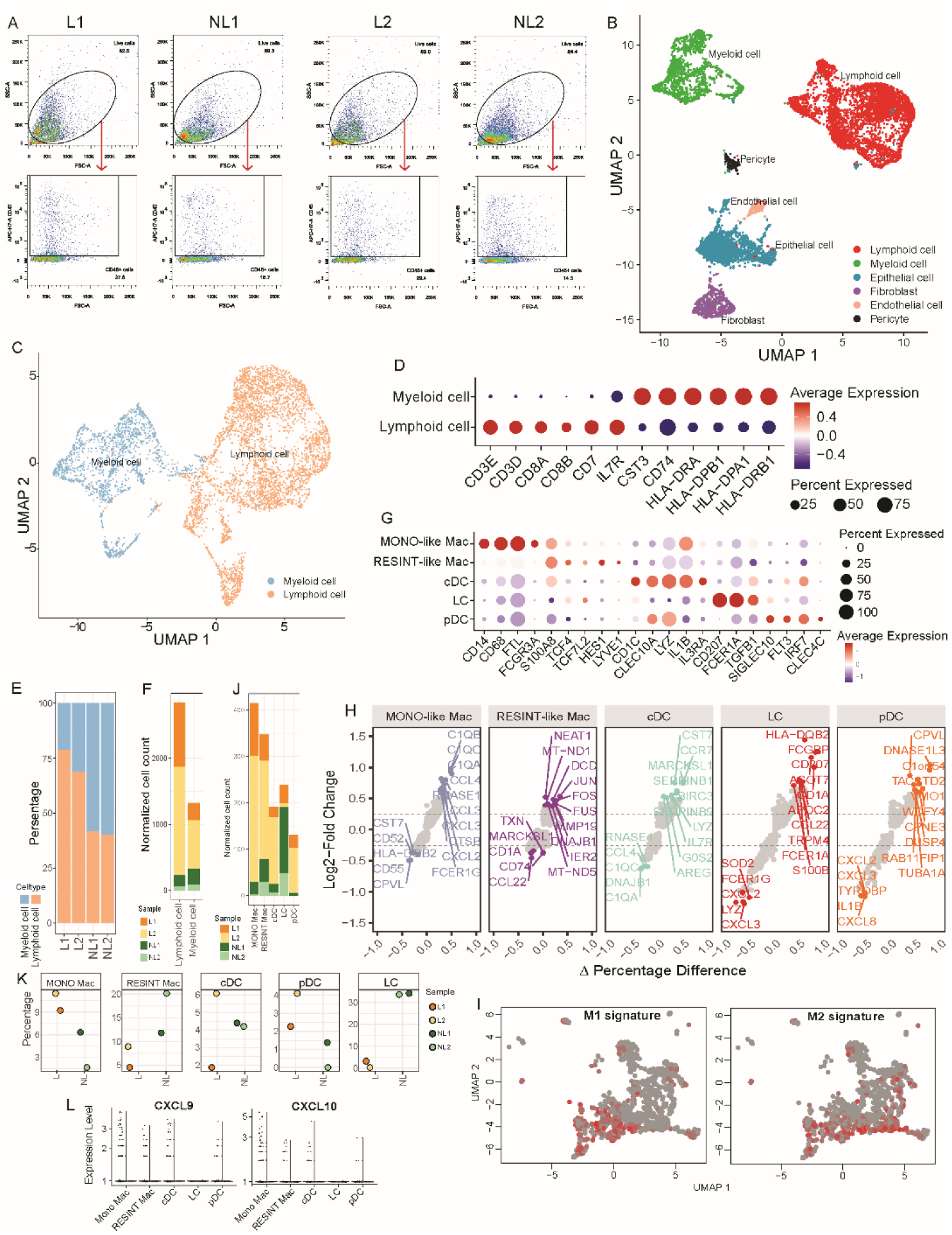
Analysis of skin myeloid cells. A. Gating strategy for sorting CD45^+^ cells from skin cell suspensions of two pairs of skin biopsies. B. UMAP visualization of total CD45^+^ cells. C. UMAP visualization of immune cells in skin. The annotations for cell clusters are based on the expression of marker genes (Figure.S6D). D. Dot plot showing expression of marker genes of myeloid and lymphoid cell subsets. Dot size indicates fraction of expressing cells, colored according to z-score normalized expression levels. E. Bar charts showing the proportions of myeloid and lymphoid cells in lesions (L) and non-lesions (NL). F. Bar charts showing the normalized cell counts (absolute cell counts were normalized by the cell number in the cell suspension of each sample) of myeloid and lymphoid cells in lesions and non-lesions. G. Dot plot showing expression of marker genes of skin myeloid cell subsets. Dot size indicates fraction of expressing cells, colored according to z-score normalized expression levels. H. Expression of signature genes of skin myeloid cell subsets. Log2fold changes between cell subset of interest and all other cells. Percentage difference = (proportion of cells expressing the gene of interest in the cell subset of interest) – (proportion of cells expressing the gene of interest in all other cells).I. M1/M2 signature of macrophages. Red dots denote cells passing the threshold of M1/M2 classification. J. Bar charts showing the normalized cell counts of myeloid cell subsets in lesions (L) and non-lesions (NL). K. Abundance of myeloid cell subsets in lesions and non-lesions. L. Expression profiles of the IFN-γ-inducible ligands CXCL9 and CXCL10 across skin myeloid cell subsets between conditions.

**Supplementary Figure 9.**
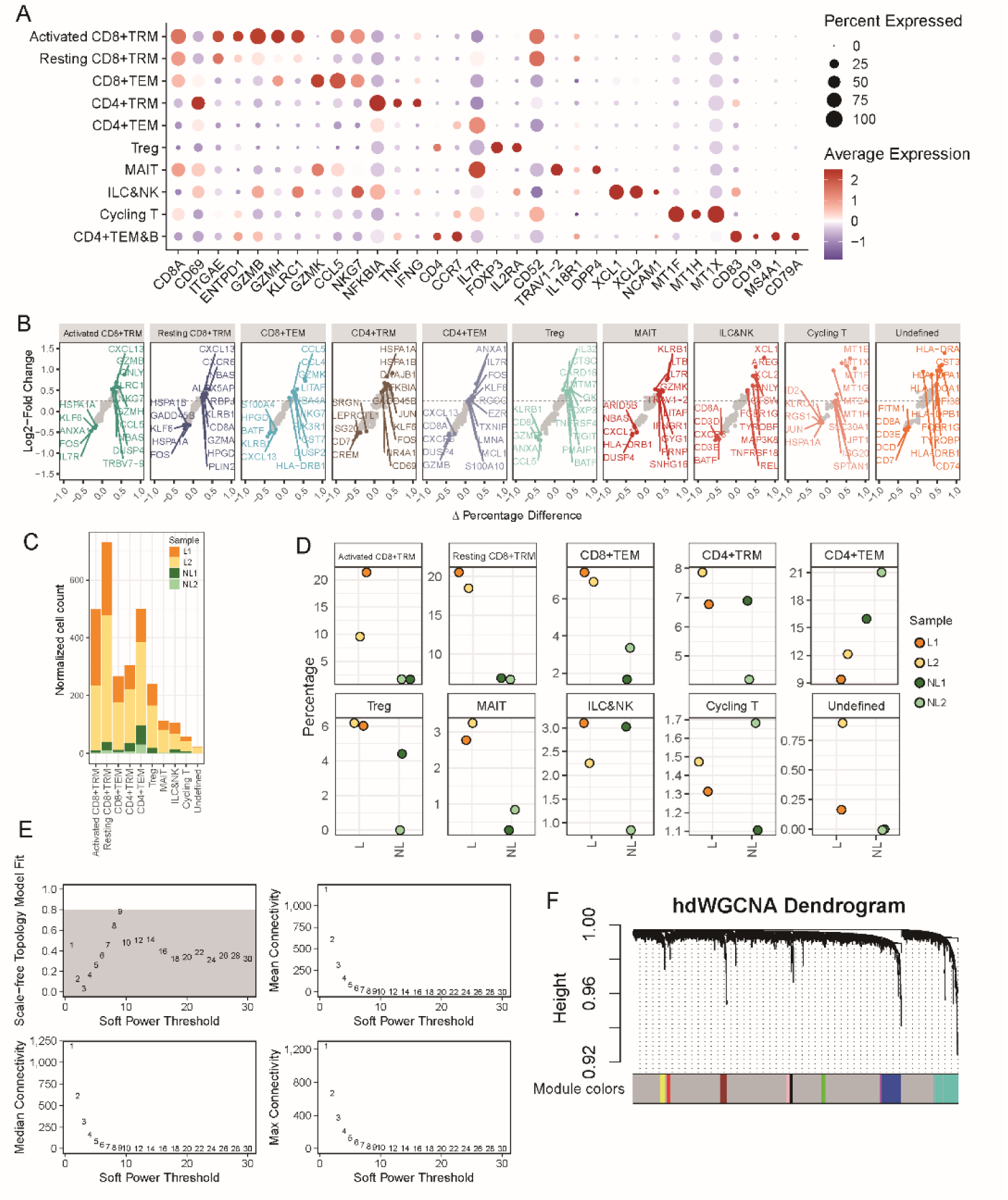
Analysis of skin lymphoid cells. A. Dot plot showing expression of marker genes of T and NK cell subsets in skin. Dot size indicates fraction of expressing cells, colored according to z-score normalized expression levels. B. Expression of signature genes of the T and NK cell subsets in skin. Log2fold changes between cell subset of interest and all other cells. Percentage difference = (proportion of cells expressing the gene of interest in the cell subset of interest) – (proportion of cells expressing the gene of interest in all other cells). C. Bar charts showing normalized cell counts of lymphoid cell subsets in lesions and non-lesions. D. Abundance of lymphoid cell subsets between conditions. Two-sided Wilcoxon test: *P<0.05. E. Different values for the soft-power threshold β were tested before co-expression network computation and gene module detection. F. Construction of the gene co-expression network.

